# Relyophilized collagen scaffold to improve handling and small molecule loading

**DOI:** 10.64898/2026.02.13.705816

**Authors:** Alison C. Nunes, Julia Harrer, Sreedhara Sangadala, Thanh A. Doan, Scott D. Boden, Nick J. Willett, Brendan A.C. Harley

## Abstract

Tissue engineering scaffolds such as collagen-based biomaterials have long been used to mimic native extracellular matrix in a wide range of regenerative applications. Their high porosity, tunable degradation and mechanics, and cell adhesion sites provide a structure upon which cells can grow and differentiate, while they also have the potential to act as carriers for loading and release of biomolecules to aid in healing. Here we describe the inclusion of a second lyophilization step in the fabrication process to enable improved loading efficiency of bone morphogenic protein 2 as well as increased ease of end-user handling. We report mineralized collagen scaffolds demonstrate maintained microarchitecture and mechanical properties post-relyophilization with reduced variability in biomolecule loading. Relyophilization allows consistent loading and release profiles and suggests the potential to improve the translational potential of collagen scaffold biomaterials for regenerative medicine applications.

## 1. Introduction

The extracellular matrix (ECM) is a complex macromolecular structure composed of a fibrillar network of structural proteins including collagen, proteoglycans and glycosaminoglycans (GAGs), specialized proteins for cell adhesion (fibronectin, laminin, etc.), and other tissue-specific materials such as apatite in bone. Tissue engineering scaffolds are used extensively as three-dimensional analogs of the ECM. Collagen-based biomaterials have a history of use in the field of regenerative medicine as ECM analogs for the regeneration of skin and peripheral nerves ^1-4^, and are currently being considered for the regeneration of a range of musculoskeletal tissues^5-8^. An important element of their success is that they possess several advantageous characteristics for regenerative biomaterials, namely a highly porous structure with interconnected pores that enables rapid cellular infiltration ^9^, tunable degradation and resorption rates ^2^, surface ligands for cell adhesion, and mechanical integrity ^10^.

In addition to providing a supportive framework for cell growth and differentiation, regenerative biomaterials can be used as carriers for biomolecules that deliver essential exogenous cues to further shape regenerative outcomes^11^. These biologic stimuli play a notable role in a number of clinically relevant tissue regeneration products, such as the use of bone morphogenic proteins (BMPs) to accelerate osteogenesis for craniofacial and extremity bone healing and regeneration^12,13^. Our lab has developed both non-mineralized collagen-glycosaminoglycan (CG)^14-16^ and mineralized collagen-glycosaminoglycan (MC)^17-19^ scaffold variants for a range of connective tissue and craniofacial bone regeneration applications. The MC scaffold variant in particular has been shown to enhance osteoprogenitor osteogenic differentiation and craniofacial bone regeneration in the absence of traditional osteogenic supplements (e.g., BMP-2) ^20^. While the lyophilization-based fabrication scheme provides opportunities to adjust pore size and shape of these scaffolds that can directly affect cell bioactivity ^1,9,21-23^, post-fabrication processing can be equally consequential in the context of ease-of-use and clinical practicality.

Carbodiimide-based crosslinking has been long used to increase crosslinking levels after fabrication, both decreasing degradation rate^2,24-26^ and increasing scaffold mechanical stiffness^27^. We showed that tuning MC scaffold mechanics significantly influenced osteoprogenitor osteogenic activity via effects on osteoprotegerin (OPG) expression and osteogenic differentiation ^28,29^. While an important processing step, carbodiimide-based crosslinking requires hydration ^30^, reducing the ease of handling, transport, and storage (hence translational potential) of a sterile, dry, sponge-like material originally formed during lyophilization.

In addition to mechanics, tissue engineering approaches often consider delivery of exogenous factors (e.g., cytokines) to promote implant integration, cellular recruitment, and regenerative activity. However use of supraphysiological doses of exogenously added growth factors^31^ can induce significant complications such as ectopic bone formation/resorption in the context of craniofacial bone repair and can significantly increase cost ^32-36^. Our lab has previously reported a number of methods for incorporating growth factors into collagen scaffolds to reduce the need for bolus dosages including covalent ^37,38^ or light-driven ^39^ immobilization to the scaffold microstructure, adjusting GAG chemistry to transiently sequester molecules via charge-charge interactions ^16,40-42^, and cyclic exposure of mineralized scaffolds to modified simulated body fluid (mSBF) to generate mineral coatings that entrap growth factor ^43^. While previous methods achieved high retention for small quantities of growth factors, all were based on adding the factors to already crosslinked (hence hydrated) scaffolds, making it difficult to calculate and control initial loading. Thus, there is a need for a method to fabricate and incorporate growth factors into a porous, crosslinked, yet non-hydrated collagen scaffold while preserving the material’s mechanical properties, sequestration capacity, and controlled release kinetics.

In this work we present a novel processing approach to enhance collagen scaffold translational potential by minimizing end-stage handling. We describe a relyophilization process that converts hydrated, crosslinked scaffolds back to dry, porous scaffolds without significant disruption of the scaffold architecture. Scaffolds are hydrated, carbodiimide crosslinked, snap-frozen, then lyophilized a second time to remove the aqueous content. We report the use of rapid freezing to cold temperatures that result in small ice crystals to reduce architectural changes induced by large ice-crystals, We assess the degree to which mineralized and non-mineralized collagen scaffolds retain their shape, structural integrity, mechanics, and porosity after relyophilization. We subsequently examine the degree to which relyophilized scaffolds exhibit improved small-molecule loading consistency and retention during loading compared to traditional scaffolds while maintaining comparable release kinetics needed to modulate cell activity.

## 2. Materials and Methods

### 2.1. Fabrication of collagen-glycosaminoglycan scaffolds

All scaffolds were fabricated by lyophilization from precursor suspensions. The mineralized collagen suspension was formed by homogenizing 1.9% w/v type I collagen (Regenity, Paramus, New Jersey USA), calcium hydroxide (Sigma Aldrich, St. Louis, Missouri USA), calcium nitrate (Sigma Aldrich), and 0.84% w/v chondroitin-6-sulfate (Sigma Aldrich) in a buffer solution containing 0.1456M phosphoric acid and 0.037 M calcium hydroxide ^6,27,43,44^. The non-mineralized collagen suspension was formed by homogenizing 0.5% w/v type I collagen and 0.05% w/v chondroitin-6-sulfate in a buffer solution of 0.05M acetic acid ^27^. The suspensions were stored at 4°C then degassed prior to lyophilization using a Genesis freeze-dryer (VirTis, Gardener, New York USA). MC and CG scaffold sheets were fabricated by pipetting 67.4mL into custom stainless-steel molds (5 in. x 5 in.) before cooling the suspensions from 20°C to - 10°C at a constant rate of 1°C/min. with a 2 hour hold at −10°C. The suspensions were subsequently sublimated at 0.2 Torr and 0°C to create a porous scaffold network. Post-fabrication, scaffolds were cut using 8mm biopsy punches (8mm diameter and 3mm height or 2mm height for MC and CG scaffolds respectively).

### 2.2. Hydration and crosslinking of scaffolds

Dry MC and CG scaffolds were hydrated and crosslinked using previously described carbodiimide crosslinking chemistry ^43,45^. Briefly, scaffolds were first hydrated for 2 hours in ethanol at ∼20°C (room temperature) under moderate shaking, then washed for 1 hour in PBS at room temperature followed by additional 30-minute and 15-minute PBS washes. A solution of 1-ethyl-3-(3-dimethylaminopropyl) carbodiimide (EDC, Sigma Aldrich Chemical Co.) and N-hydroxysuccinimide (NHS, Sigma Aldrich Chemical Co.) at a 5:2:1 ratio of EDAC:NHS:COOH (COOH groups are the collagen reactive sites) was used to crosslink the scaffolds for a total of 1.5 hours at room temperature. Following this, scaffolds underwent sequential 45-minute and 15-minute PBS washes.

### 2.3. Relyophilization of collagen-glycosaminoglycan scaffolds

After hydration and crosslinking scaffolds, a second lyophilization step was used to create relyophilized scaffolds. Briefly, hydrated 8mm scaffold disks were transferred using tweezers into 48-well plates to provide structural support. The well-plates were then placed at −80 °C in a Isotemp Ultra Low Temperature Freezer (Thermo Fisher Scientific, Waltham, MA) to rapidly freeze the hydrated scaffolds to minimize the formation of large ice crystals^46^ that would disrupt the original pore architecture. Well-plates were then transferred to the lyophilizer with a pre-cooled shelf temperature of −40°C for an additional 2-hour hold. Frozen scaffolds were subsequently sublimated at 0.2 Torr and 0°C to recover the porous scaffold network.

### 2.4. Gross morphology and mechanical testing of mineralized and non-mineralized collagen scaffolds

Calipers were used to measure the diameter and height of the scaffolds in triplicate. Unconfined, unidirectional compression tests were performed on dry scaffolds (mineralized and non-mineralized) for regular and relyophilized conditions (n=6) as previously described ^47^. Briefly, tests were performed using an Instron 5943 mechanical tester (Instron, Norwood, Massachusetts USA) using a 5N load cell at a constant strain rate (0.1% strain s-1) perpendicular to the plane of the scaffold sheet. Stress-strain curves were generated, and the Young’s modulus was determined using a custom MATLAB code using conventional analysis methods for low-density open-cell foam structures ^27,48,49^. Mechanical properties were reported for dry samples only comparing the effects of relyophilization (previous work descried the role of hydration and EDC-NHS crosslinking on scaffold mechanics^27,29,49^).

### 2.5. Quantifying calcium and phosphorous deposition in mineralized collagen scaffolds

The mineral content of MC scaffolds and relyophilized MC scaffolds was analyzed via inductively coupled plasma emissions spectrometry (ICP-OES) ^50,51^. Dry samples (n=8) were transferred to a digestion tube containing concentrated nitric acid (67-70%) before undergoing automated sequential microwave digestions using a CEM Mars 6 microwave digester (CEM Microwave Technology Ltd., North Carolina USA). The resulting clear aqueous solution was diluted with DI water to a total volume of 25mL before being run through an inductively coupled plasma-mass spectrometer (NexION^TM^ 350D ICP-MS, PerkinElmer, USA).

### 2.6. Micro-CT images

Scaffold parameters were measured using a MicroXCT-400 (Zeiss, Oberkochen, Germany) with n=3 scaffolds scanned per group. Scans utilized a 20x objective with a field of view size of 808.57μm, voltage of 100kV and voxel size of 0.81μm. To quantify pore architecture and strut structure, the image processing and data analysis software Dragonfly (Comet Technologies Canada Inc, Montreal, Canada) was used.

### 2.7. Scanning electron microscopy of original and relyophilized scaffolds

Scanning electron microscopy was used to qualitatively compare the pore structure of lyophilized and relyophilized scaffolds ^8^. Dry scaffolds (n=2) were cut to expose the interior architecture before sputter coating with Au/Pd (Denton Desk II TSC, New Jersey, USA). Samples were then imaged using an FEI Quanta FEG 450 ESEM (FEI, Hillsboro, OR) under high vacuum.

### 2.8. Loading and elution of bone morphogenetic protein 2 onto collagen scaffolds

Bone morphogenetic protein 2 (rhBMP-2) was diluted in 1% BSA in PBS to generate 40μg/mL or 200μg/mL solutions. 25µL of the rhBMP-2 solution was subsequently added to either hydrated MC scaffolds (After EDC crosslinking) or dry relyophilized MC scaffolds in individual wells of a 24-well plate (loading either 2μg or 10μg rhBMP-2, respectively). After 3 minutes, scaffolds were transferred into new wells containing fresh PBS at 37 °C. Aliquots were collected (and replaced) at 13 timepoints to measure rhBMP-2 release: 1 min., 5 min., 10 min., 15 min., 30 min., 60min., 2hrs., 4hrs., 6hrs., 12hrs., 24 hrs., 48 hrs., 72hrs. Collected samples were stored at −20°C prior to analysis. Separately, 1mL of PBS was added to the original loading wells to collect any loading solution that was not retained in the scaffold; these samples were also stored at −20°C prior to analysis.

To account for all rhBMP-2 originally loaded into scaffold, a 6-hour elution study was followed by specimens being transferred to new wells containing 1mL of 1M NaCl to elute any rhBMP-2 ionically bound to the scaffold. Scaffolds sat in the elution buffer overnight before the 1M NaCl was collected and replaced with fresh elution buffer for an additional overnight soak. All collected buffer was stored at −20°C prior to analysis.

### 2.9. Quantifying rhBMP-2 release from MC and relyophilized MC scaffolds

An enzyme-linked immunosorbent assay (ELISA) (R&D Systems, Minneapolis, MN, USA) was used to quantify the amount of rhBMP-2 loaded and released from each scaffold groups over time per previously reported protocols ^52^. The assay was conducted according to the manufacturers protocol with a 10x sample dilution and PBS as a background control. Data was reported as cumulative release of rhBMP-2 from each scaffold group.

### 2.10. Statistics

Statistics were performed using RStudio (RStudio, Massachusetts, USA) and OriginPro (OriginPro, Massachusetts, USA). “n” values in methods indicate number of technical replicates run. Significance was set to p<0.05. A Shapiro-Wilks test was used to test for normality followed by a Grubbs test to remove outliers if data was not normal. If an outlier was identified, but removal did not result in normal data, the outlier was not removed from the sample set. A Levene’s test was then performed to test the equal variance assumption. For normal data sets with equal variance, an ANOVA with a Tukey post-hoc test was run. For normal data sets with unequal variance, a one-way Welch’s ANOVA and Welch/Games-Howell post-hoc was performed. For non-normal data sets with unequal variance, a Welch’s Heteroscedastic F test and Welch/Games-Howell post-hoc were performed. Finally, for non-normal data with equal variance, a Kruskal-Wallis test was performed. Error bars represent mean ± standard deviation, and all graphs were created in OriginPro.

## 3. Results

### 3.1. Relyophilized scaffolds do not display significantly altered gross structural or mechanical properties

We compared the *dry/relyophilized* and *(re)hydrated/relyophilized* properties of relyophilized scaffolds to (1) *dry/uncrosslinked* or (2) *hydrated/crosslinked* original scaffold specimens (**Figure 1**). *Dry/relyophilized* (crosslinked) scaffolds maintained an open pore network as well as similar size and morphology as *dry/uncrosslinked* variants **(MC: Figure 1, 2A/B/C; CG: Figure S1)**. ICP analysis of MC scaffolds confirmed that relyophilization did not alter the relative amount of calcium or phosphate in the scaffold architecture (**Figure 2D**). Lastly, we observed no difference in the Young’s modulus of non-crosslinked MC scaffolds before or after relyophilization (**Figure 2E**); it is not possible to have dry/crosslinked scaffolds in the original (non-relyophilized) format for comparison with *crosslinked/relyophilized* variants.

**Figure 1.**
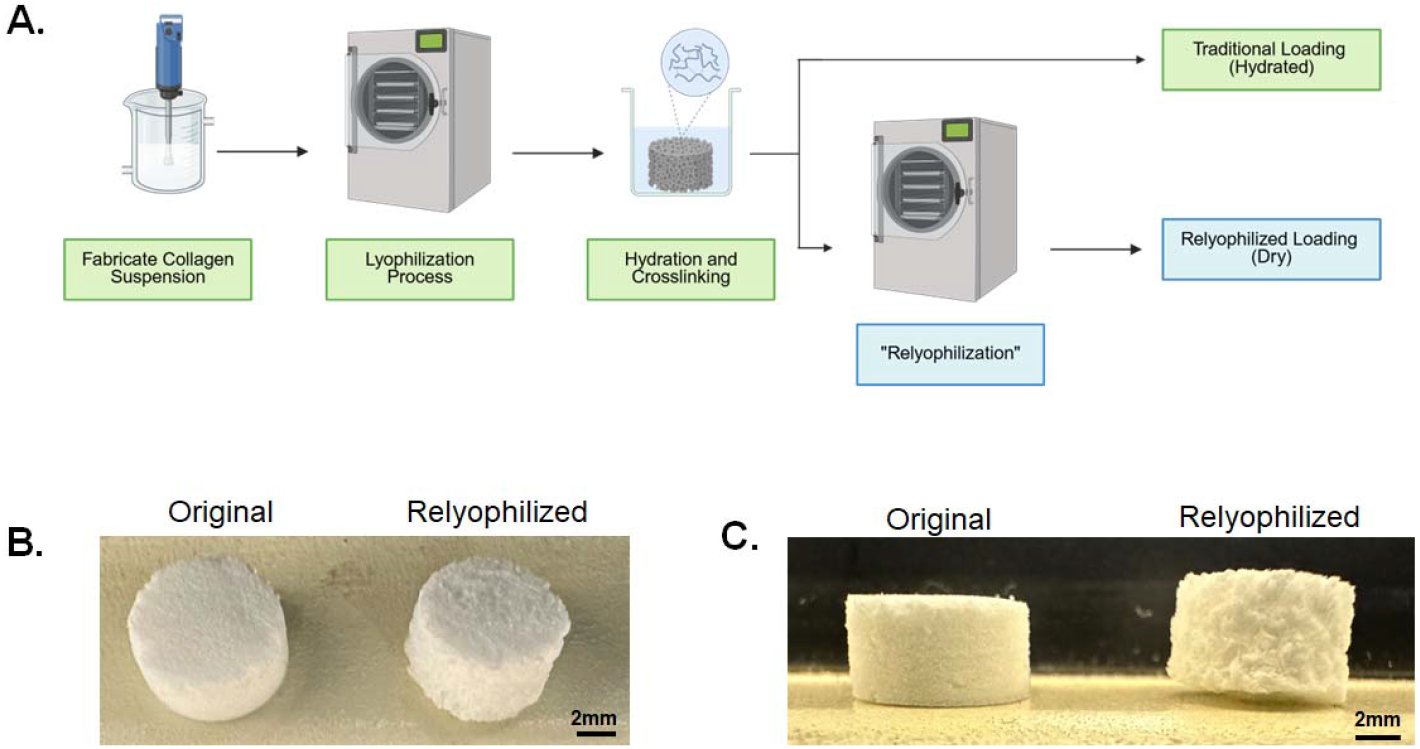
(a) Collagen scaffolds underwent a hydration and crosslinking process. Traditional loading techniques utilized direct pipetting onto hydrated scaffolds, while the relyophilized scaffolds underwent an additional lyophilization step such that loading occurred via direct pipetting onto dry scaffolds. (b) Top view of 8mm MC scaffold and relyophilized MC scaffold from left to right respectively. (c) Side view of 8mm MC original scaffold and relyophilized MC scaffold from left to right respectively.

**Figure 2.**
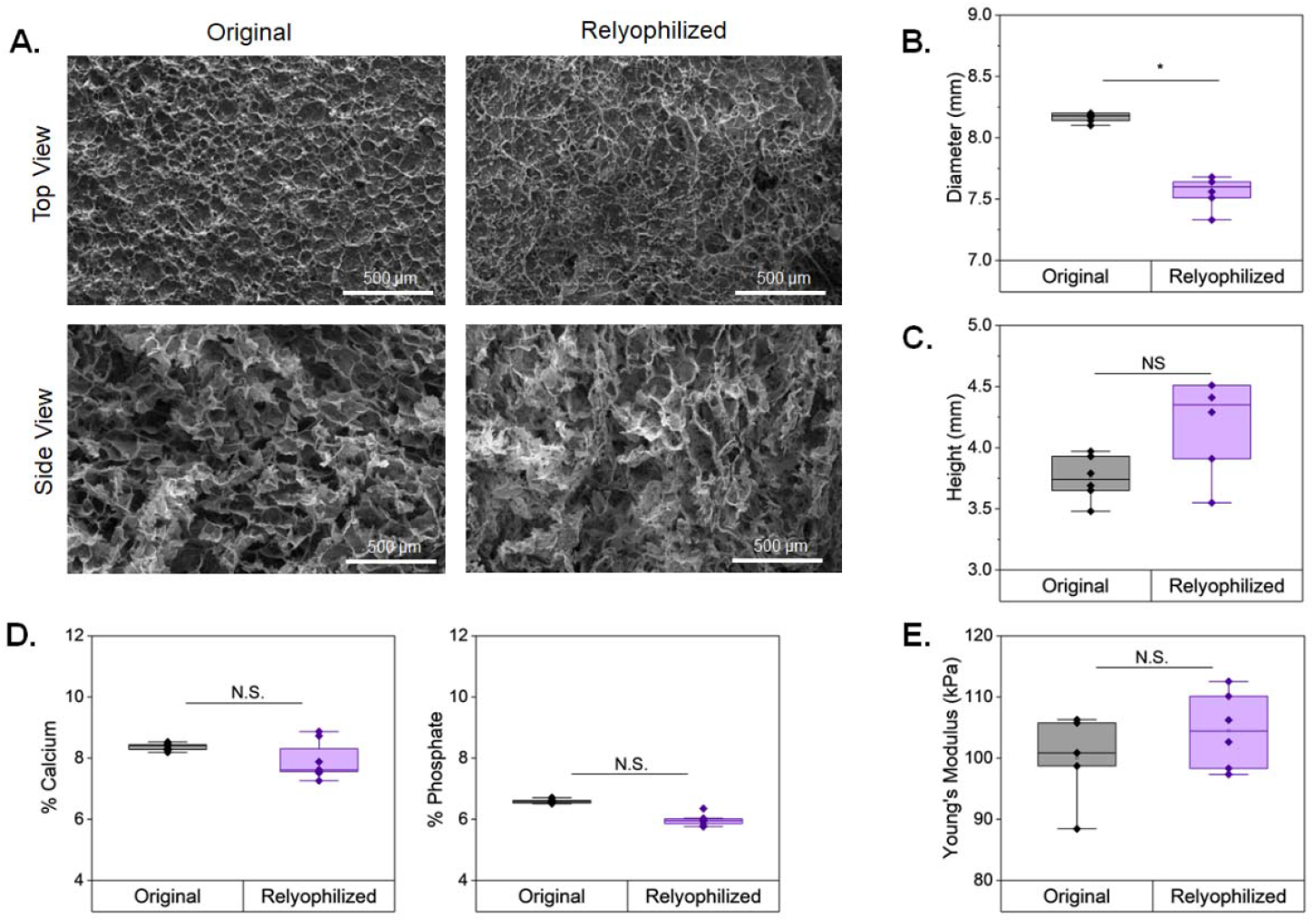
Physical characterization of traditional mineralized collagen scaffolds before and after undergoing hydration, crosslinking, and second lyophilization step to become relyophilized mineralized scaffolds. (a) eSEM images (100x) showing top views and side views of the pore architecture for mineralized original and relyophilized scaffolds. (b-c) Changes in diameter and height (mm) of mineralized scaffolds pre- and post-relyophilization with a significant change in diameter (p<0.05) but no significant change in height. (d) Percent of calcium and phosphate contained in original and relyophilized scaffolds with no significant change in either. (e) Young’s modulus before and after relyophilization shows no significant increase.

### 3.2. Relyophilization does not affect the microstructural organization of MC scaffolds

We subsequently performed µCT analysis of (non-mineralized) CG and (mineralized) MC scaffold variants to assess scaffold microarchitecture at a resolution (0.81μm voxel size) sufficient to examine scaffold strut morphology. MC scaffolds remained highly porous after crosslinking and relyophilization (**Figure 3A**), with no significant changes in metrics of scaffold porosity, average pore size, and average strut thickness between (original) non-crosslinked scaffolds and relyophilized/crosslinked scaffolds (**Figure 3B/C/D**). CG scaffolds showed a slight but significant reduction in overall porosity (p<0.01) and increase in average strut thickness (p<0.01) after crosslinking and relyophilization versus scaffolds as fabricated, but no overall change in the mean pore size (**Figure 4**).

**Figure 3.**
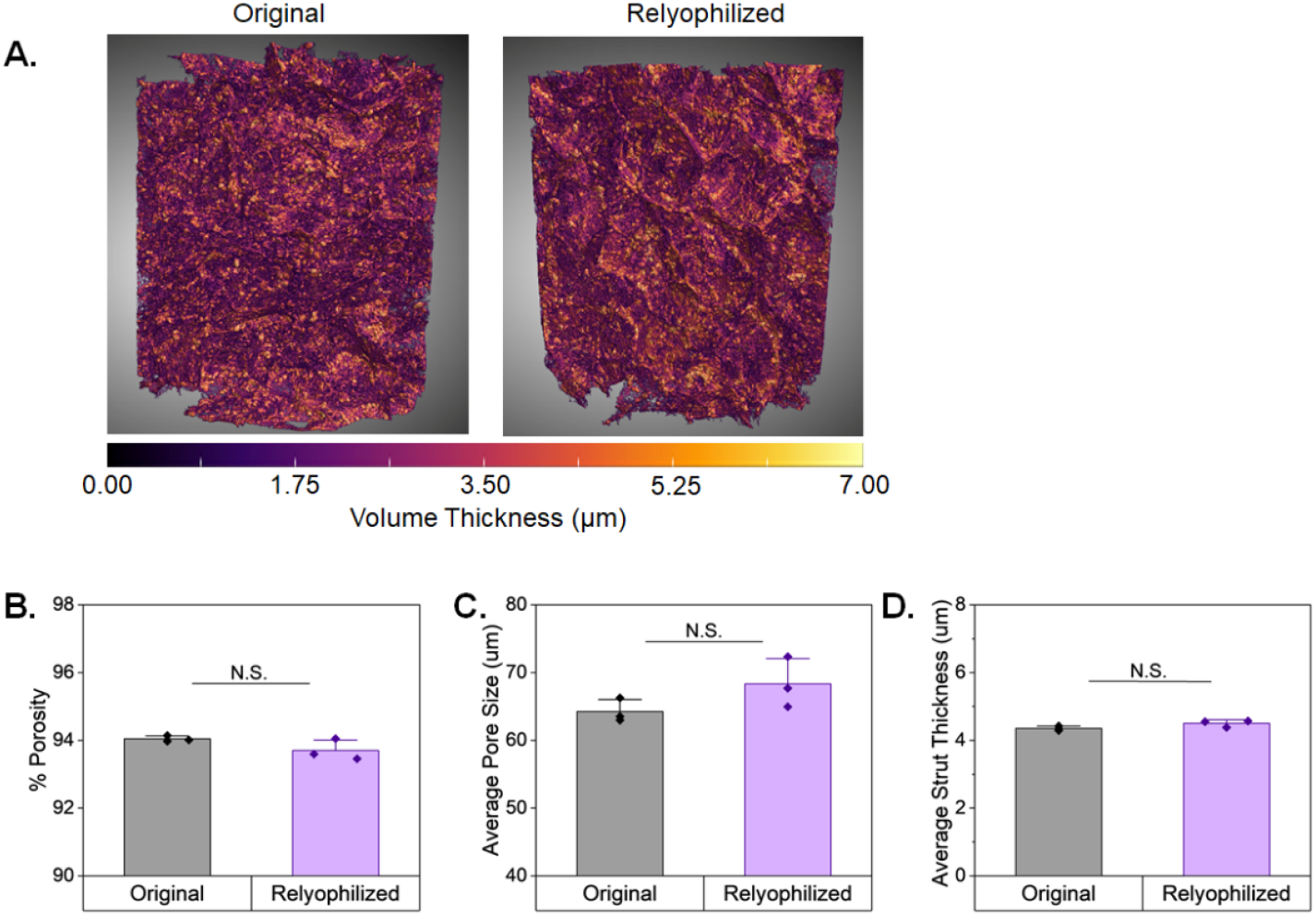
(a) Representative 3D reconstruction of uCT images comparing the volume thickness of pores in original and relyophilized mineralized collagen scaffolds. (b) Percent porosity analysis from uCT data. (c) Average pore size as calculated from uCT data. (d) Average strut thickness as calculated from uCT data for mineralized collagen scaffolds. There is no significant difference in the pores or struts of original compared to relyophilized mineralized scaffolds.

**Figure 4.**
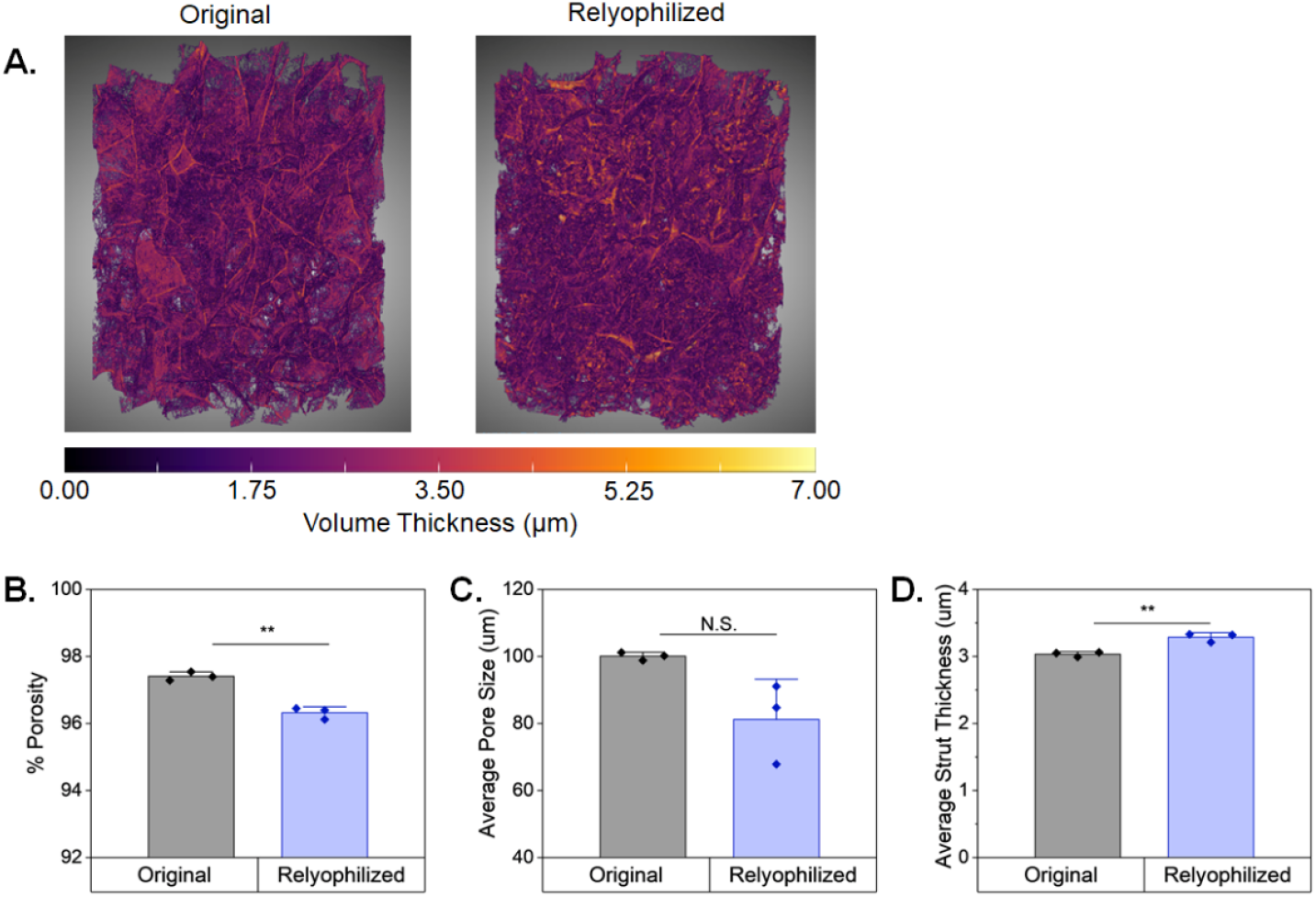
(a) Representative 3D reconstruction of uCT images comparing the volume thickness of pores in our original and relyophilized non-mineralized collagen scaffolds. (b) Percent porosity analysis from uCT data showed significant (p<0.01) reduction in porosity post-relyophilization. (c) Average pore size as calculated from uCT data showed no significant differences. (d) Average strut thickness as calculated from uCT data for non-mineralized collagen scaffolds and showed significant (p<0.01) increase in strut thickness post-relyophilization.

### 3.3. Relyophilized scaffolds provide a more consistent platform for small molecule loading and release

We subsequently examined changes in rhBMP-2 incorporation and release due to relyophilization **(Figure 5A)**. A significant concern for biomolecule loading in traditional (non-relyophilized) scaffolds is that the required crosslinking process leaves a hydrated scaffold to which a growth factor solution is added. The unknown initial volume of aqueous solution trapped in the hydrated scaffold complicates calculating dilutions factors for release studies and prediction of how rapidly biomolecules in a solution penetrate and adhere to the scaffold struts. However, a small loading volume can be quickly and completely wicked into a dry (e.g. relyophilized) scaffold. Hence, scaffold relyophilization provides an entirely new way for small molecule loading. Relyophilized MC scaffolds showed significant improvement (p<0.05) in small molecule retention compared to original (hydrated, non-relyophilized) MC scaffolds for both low (2μg) and high (10μg) rhBMP-2 **(Figure 5B)**. Additionally, for the low initial loading (2μg) group there was no significant difference in the release of rhBMP-2 over the first 6-hours **(Figure 5C)**. However, when scaled up to a high (10μg) initial loading condition, relyophilized scaffolds showed a significant reduction (p<0.05) in rhBMP-2 released starting a the 2-hour mark **(Figure 5D)**. For CG scaffolds, there were no significant differences in loading efficiency between original or relyophilized scaffolds for either loading group (low (2μg) or high (10μg)). **(Figure S2A)**. The release profile of CG scaffolds loaded with low doses of rhBMP-2 showed a significant (p<0.01) decrease in rhBMP-2 released from relyophilized scaffolds compared to original, while the high loading groups showed no significant differences in release between relyophilized and original CG scaffolds **(Figure S2B/C)**.

**Figure 5.**
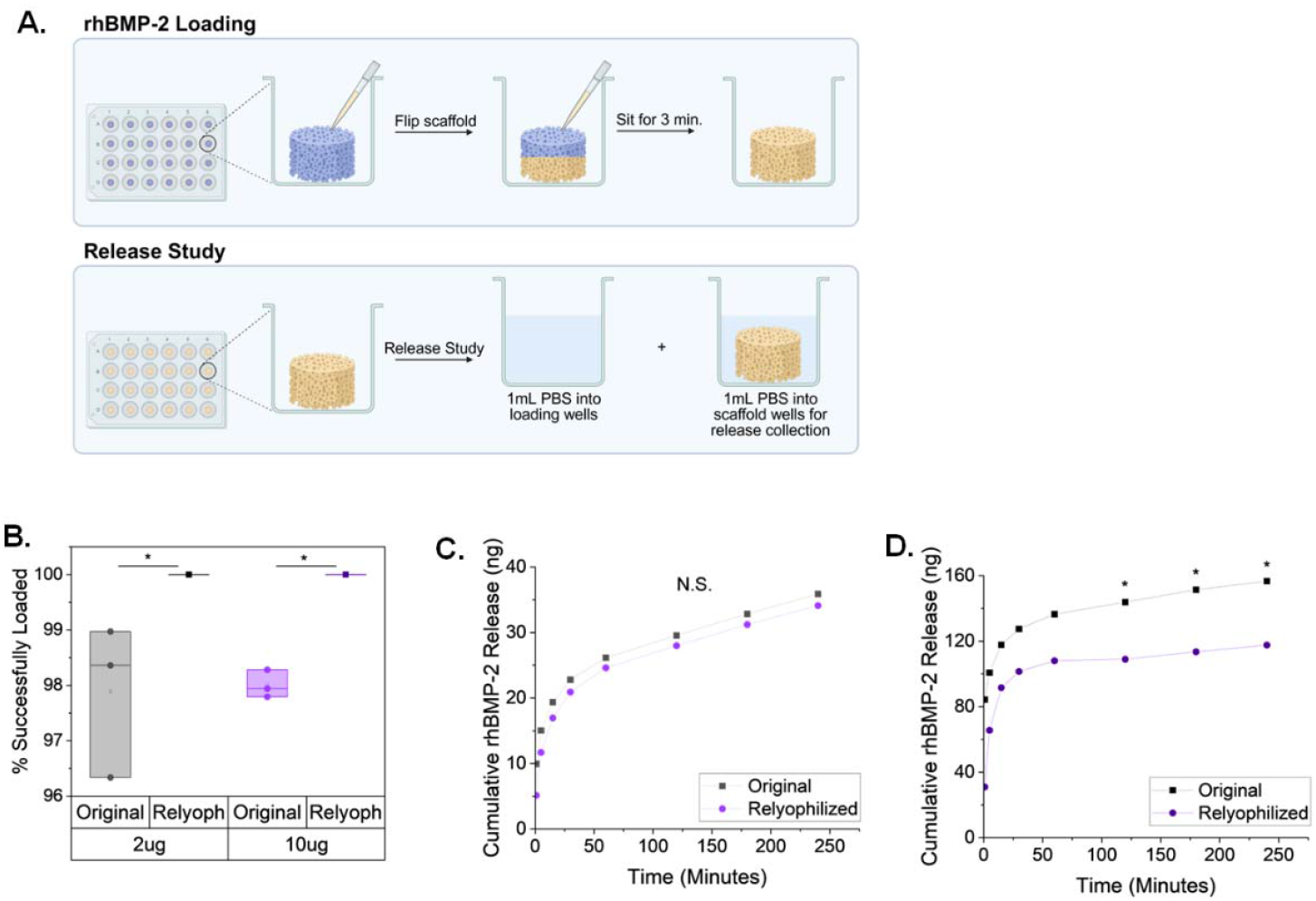
(a) Collagen scaffolds were placed in a 24-well plate where half the volume of rhBMP-2 was loaded before scaffolds were flipped and the remaining volume of rhBMP2 was direct pipetted onto the scaffold. Scaffolds sat for 3 minutes before transfer into new 24-well plates for release study. Original scaffolds were loaded in a hydrated state, while relyophilized scaffolds were loaded in a dry state. 1mL of PBS was added to the loading well post-loading to collect any rhBMP-2 that was unsuccessfully loaded onto the scaffolds. (b) Relyophilized mineralized scaffolds showed 100% loading efficiency for both 2ug and 10ug loading conditions showing significant (P<0.05) increase from original scaffolds. (c) Cumulative release of rhBMP-2 from original and relyophilized scaffolds showed no-significance out to 6hrs for the 2ug initial loading condition, while the 10ug loading condition (d) showed significant decrease (p<0.05) in rhBMP-2 release from 2hours onward.

## 4. Discussion

Collagen scaffolds have been widely deployed for regenerative medicine applications. Non-mineralized versions have seen clinical translation for skin and peripheral nerve regeneration^2,4,53-55^ while mineralized and gradient (mineralized-nonmineralized) versions are under preclinical development for a range of musculoskeletal (*e*.*g*., rotator cuff, osteochondral, extremity bone, craniofacial bone) applications ^5,22,31,56-59^. These scaffolds have been traditionally deployed in an acellular format, relying on the recruitment of cells from the wound periphery to enable regenerative healing. Versions of this scaffold have also been optimized to induce the endogenous production of activity-inducing cytokines; notably, the mineralized scaffold variant studies here have been previously optimized to increase osteoprogenitor (e.g., mesenchymal stem cell, MSC) production of osteoprotegerin^7,60-63^, a glycoprotein that can transiently inhibit osteoclast activity and aid craniofacial bone regeneration. However, indogenous production of cytokines is often not sufficient to enable functional regenerative healing in large^64^ compared to more moderate^17^ sized defects. As a result, prior work has also explored strategies to deliver biomolecules from the scaffold, notably via transient sequestration via incorporation of proteoglycans in the scaffold^15,16^ or via other non-covalent means ^52^.

One challenge in employing these collagen scaffolds for clinical applications is their extensive end-user handling. Scaffolds undergo a hydration and carbodiide crosslinking process with a total minimum time of 6.5 hours to get the material is ready for implantation, all of which must be conducted in a biosafety cabinet and aseptic technique after ethylene oxide sterilization of the scaffolds which must take place on the dry scaffold^7^. TRelyophilizationallows for hydration and crosslinking of the collagen scaffold material prior to sterilization which can now be applied to the relyophilized, crosslinked product, allowing for more flexibility at the time of implantation to assess wound geometry and cut the (dry) material to size. Another challenge with the conventional collagen scaffolds is a poor understanding of initial loading of exogenous factors. Due to their hydrated state and difficulty in controlling the exact degree of hydration in each scaffold specimen, it is difficult to quantify exact loading of exogenous factors into the already hydrated scaffold. The development of a dry, relyophilzied collagen scaffold allows for controlled loading of factor in a smaller dilution volume that can be completely absorbed by the scaffold.

In this study we hypothesized that the addition of a second lyophilization step after material hydration and crosslinking would result in more accurate exogenous factor loading with reduced sample to sample variability, as well as a simplification of end-user handling to improve translational potential. We employed a series of material and mechanical characterization techniques to confirm consistent structural and mechanical properties post-relyophilization. A primary concern was that freezing a hydrated scaffold would result in large icre crystals that disrupt the scaffold architecture. Prior studies of scaffold lyophilization showed that decreasing freezing temperature and increasing speed of freezing result in smaller ice crystals in the aqueous phase^1,3^. Here, scaffolds were fabricated via a slow temperature ramp to −10°C followed by an extended hold to increase ice crystal coarsening^6^ and ultimate pore size, but hydrated scaffolds were then relyophilized via rapid freezing to −80°C. Analysis of scaffold microarchitecture via micro-CT showed that relyophilization does not significantly change the strut thickness, porosity, or average pore size. The dependence on the mechanical performance of low-density open-cell foams on their porosity has established that even subtle changes in foam architecture can result in significant changes in mechanics^27,65,66^. Mechanical testing confirmed no changes in the Young’s modulus of lyophilized MC scaffolds, further confirming material and mechanical properties of the MC scaffolds are maintained after relyophilization. Interestingly, non-mineralized collagen scaffolds showed a 3-fold increase in modulus consistent with the observed small but significant reduction in scaffold height and diameter as well as porosity (∼97% to ∼96%). Interestingly, the modulus of porous materials such as these scaffolds is known to scales as material relative density (1 – percent porosity) squared^27,65,66^; the increase in modulus of relyophilized non-mineralized scaffolds is consistent with a ∼60% increase in scaffold relative density, which is consistent with our observations (∼2.5% to 3.7% relative density). This suggest relyophilization may be particular valuable for mineralized collagen scaffolds where the mineral content and increased matrix density created scaffold structural features that better resist relyophilization-mediated densification. Finally, we looked at whether relyophilization influences the rate of release of rh-BMP2 from the scaffold constructs as well as the initial loading efficiency. Reylophilized MC scaffolds demonstrated improved loading compared to original (hydrated) scaffolds, and both mineralized and non-mineralized relyophilized scaffolds showed reduced variability in rh-BMP2 loading, suggesting improved consistency. We found no significant difference in BMP2 release from relyophilized versus conventional MC scaffolds for low (2μg) loading concentrations, but that relyophilied MC scaffolds displayed significantly slower release of BMP when loaded at higher concentrations (10μg). These observations are all consistent with increased consistency of loading in relyophilized scaffolds. These findings suggest future studies to examine the biological effect of increased factor retention on cell bioactivity such as improved osteogenic response for scaffolds with BMP2 embedded into the mineral content^52^; for example osteoprotengerin is a glycoprotein known to transiently reduce osteoclast activity and whose addition to MC scaffolds increases osteogenic potency^61^. Hence relyophilized MC scaffolds may provide a more consistent delivery platform for osteoprotegerin to boost scaffold regenerative potency in vivo.

## 5. Conclusions

Here we describe the impact of relyophilization on the structural, mechanical, and biomolecule loading characteristics of a class of collagen scaffolds under development for tissue regeneration applications. Mineralized collagen scaffolds demonstrate maintained microarchitecture and mechanical properties post-relyophilization with reduced variability in biomolecule loading. Relyophilization allows consistent loading and release profiles and suggests the potential to improve the translational potential of collagen scaffold biomaterials for regenerative medicine applications. These results highlight opportunities to create Relyophilized collagen biomaterials material with improved potential for clinical translation and streamlined handling, offering opportunities for improving the regenerative activity of collagen biomaterials via more consistent incorporation of biomolecules.

## Acknowledgements

The authors would like to acknowledge the following institutes for access to their facilities and services: the School of Chemical Sciences Microanalysis Laboratory, the Carl R. Woese Institute for Genomic Biology, and the Beckman Institute for Advanced Science and Technology, located at the University of Illinois. Additional support was provided by the Carl R. Woese Institute for Genomic Biology and the Chemical and Biomolecular Engineering Dept. at the University of Illinois at Urbana-Champaign. This manuscript is the result of funding in whole or in part by the National Institutes of Health (NIH). It is subject to the NIH Public Access Policy. Through acceptance of this federal funding, NIH has been given a right to make this manuscript publicly available in□PubMed Central upon the Official Date of Publication, as defined by NIH. Research reported in this publication was supported by the National Institute of Dental and Craniofacial Research of the National Institutes of Health under Award Number R01 DE030491 (BACH), National Institute of Arthritis and Musculoskeletal and Skin Diseases under Award Numbers R01 AR077858 (BACH) and R01 AR078892 (SDB/NJW). We acknowledge the use of BioRender to create Figures 1a and 5a.

## Contributions (CRediT: Contributor Roles Taxonomy ^67,68^)

**A.C. Nunes:** Conceptualization, Data curation, Formal Analysis, Visualization, Investigation, Methodology, Writing – original draft, Writing – review & editing. **J. Harrer:** Investigation, Data curation, Methodology, Formal Analysis, Writing – review. **S. Sangadala:** Investigation. **T.A. Doan:** Investigation; Project administration. **S.D. Boden:** Supervision; Funding acquisition; Writing – review & editing. **N.J. Willett:** Supervision, Funding acquisition; Methodology, Writing – review & editing. **B.A.C. Harley:** Conceptualization, Resources, Project administration, Funding acquisition, Supervision, Writing – review & editing.

## Notes

### Competing Interest Statement

The authors have declared no competing interest.

