## Supplemental Information for "Relyophilized collagen scaffold to improve handling and small molecule loading"

^2^ Dept. Chemical and Biomolecular Engineering, UIUC, Urbana, IL 61801

^3^ Department of Bioengineering, Knight Campus, University of Oregon, Eugene, OR 97403

^4^ School of Medicine, Emory University, Atlanta, GA 30322

^5^ Cancer Center at Illinois, UIUC, Urbana, IL 61801

**Corresponding Author:**

B.A.C. Harley

Dept. of Chemical and Biomolecular Engineering

Cancer Center at Illinois

Carl R. Woese Institute for Genomic Biology

University of Illinois at Urbana-Champaign

110 Roger Adams Laboratory

600 S. Mathews Ave.

Urbana, IL 61801

**Supp. Table 1:** Parameters for analysis of calcium and phosphorus collagen samples via Optima 8300 ICP-OES (see **section 2.5**). Emission lines: Ca ((II)-317.93 nm) and P ((I)- 213.62 nm).

| **ICP-OES Parameters** | **Values** |
| --- | --- |
| RF Power | 1500 Watts |
| Nebulizer | GemCone Low Flow |
| Nebulizer Gas Flow rate | 0.85L/min |
| Plasma Gas Flow rate- Argon | 10L/min |
| Sample Flow rate | 1.50mL/min |

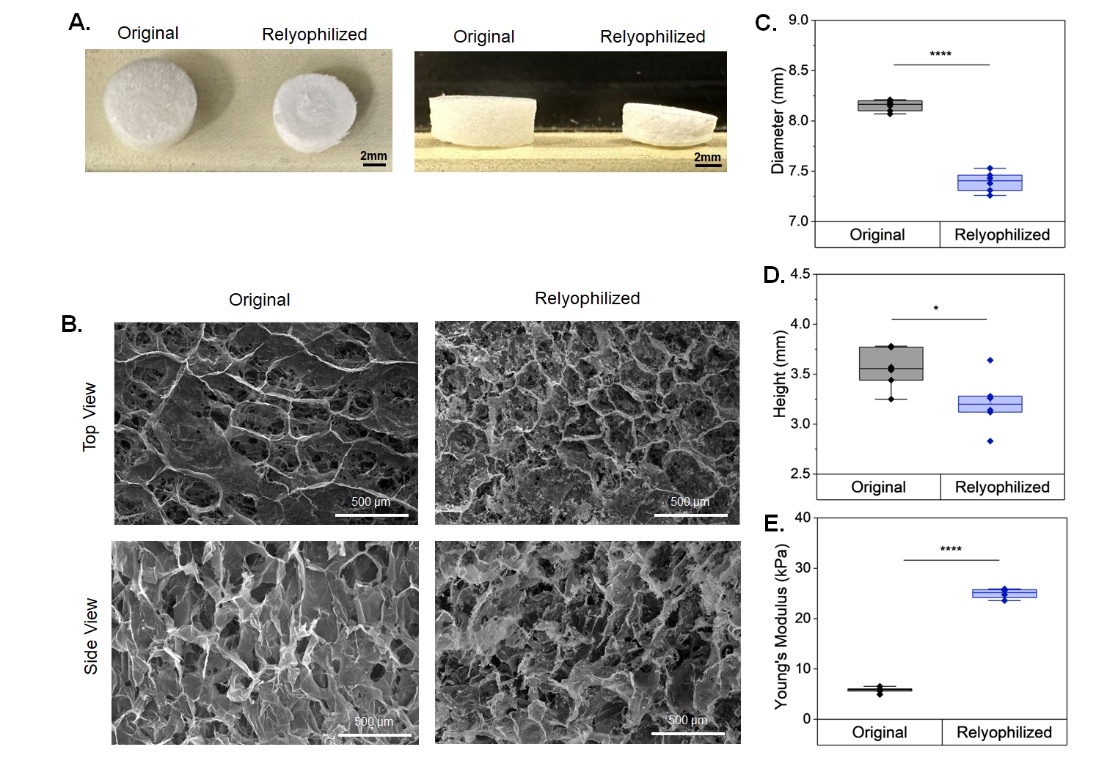

**Supplemental Figure 1.** Non-mineralized scaffolds showed significant differences in physical properties after undergoing hydration, crosslinking, and a second lyophilization step. (a) Top view of 8mm CG scaffold and relyophilized CG scaffold and side view of 8mm CG original scaffold and relyophilized CG scaffold from left to right respectively. (b) eSEM images (100x) showing top views and side views of the pore architecture for non-mineralized original and relyophilized scaffolds. (c-d) Changes in diameter and height (mm) of mineralized scaffolds pre- and post- relyophilization showed a significant (p<0.0001 and p<0.05 respectively) reduction in diameter and height. (e) Young’s modulus showed a significant (p<0.0001) increase after undergoing hydration and crosslinking.

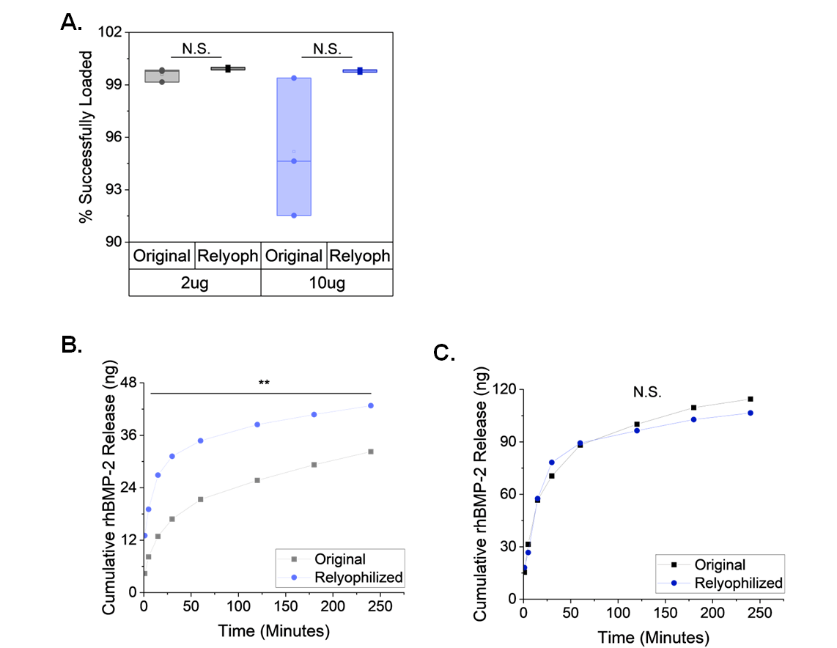

**Supplemental Figure 2.** (a) Relyophilized non-mineralized scaffolds showed almost 100% loading efficiency for both 2ug and 10ug loading conditions showing no significance from the original scaffolds, although more consistent loading efficiency is seen. (b) Cumulative release of rhBMP-2 from original and relyophilized scaffolds showed significant (p<0.01) decrease in release for the relyophilized scaffolds at all timepoints for the 2ug initial loading condition, while the 10ug loading condition (c) showed no significant difference in rhBMP-2 release.
